# A systems biology approach to explore the impact of maple tree dormancy release on sap variation and maple syrup quality

**DOI:** 10.1101/329367

**Authors:** Guillaume N’guyen, Nathalie Martin, Mani Jain, Luc Lagacé, Christian R Landry, Marie Filteau

## Abstract

Maple sap is a complex nutrient matrix collected during Spring to produce maple syrup. The characteristics of sap change over the production period and its composition directly impacts syrup quality. This variability could in part be attributed to changes in tree metabolism following dormancy release, but little is known about these changes in deciduous trees. Therefore, understanding the variation in sap composition associated with dormancy release could help pinpoint the causes of some defects in maple syrup. In particular, a defect known as “buddy”, is an increasing concern for the industry. This off-flavor appears around the time of bud burst, hence its name. To investigate sap variation related to bud burst and the buddy defect, we monitored sap variation with respect to a dormancy release index (S_bb_) and syrup quality. First, we looked at variation in amino acid content during this period. We observed a shift in amino acid relative proportions associated with dormancy release and found that most of them increase rapidly near the point of bud burst, correlating with changes in syrup quality. Second, we identified biological processes that respond to variation in maple sap by performing a competition assay using the barcoded *Saccharomyces cerevisiae* prototroph deletion collection. This untargeted approach revealed that the organic sulfur content may be responsible for the development of the buddy off-flavor, and that dormancy release is necessary for the appearance of the defect, but other factors such as microbial activity may also be contributing.

## Introduction

Maple syrup is a natural sweetener obtained from the concentration of sap collected in spring time from sugar maple trees (*Acer saccharum Marsh*.) and related species (*Acer rubrum, Acer nigrum*). The sap composition is the product of a complex system involving tree metabolism and subsequent microbial activity during the collection and transformation process, which includes storage and membrane processing, such as reverse osmosis, prior to boiling. The sap is collected through networks of plastic tubing connected to hundreds of trees under a high vacuum and stored in a non-aseptic environment, where microbial activity can alter its composition^1–4^. Most maple syrup characteristics such as color, dominant flavors, pH and chemical composition vary over the production period^5–7^. Previous studies focusing on maple sap and syrup properties have mainly considered the cumulative percentage of sap flow or syrup color grade, corresponding to its light transmittance, as a reference to monitor variation^5,8^. Although maple sap composition has been investigated^5^, its complexity and variation is such that many compounds that can contribute to its properties are still unidentified^6,9–13^.

Maple sap composition is a critical factor for syrup quality since it is concentrated 35 to 40 fold during the production process. The production of maple syrup has doubled over the last two decades, reaching 207 million pounds in 2016^14^. However, the variation of maple syrup quality has remained a constant concern, as on average 25% of the bulk production does not meet the standard quality ^14,15^. Trace off-flavors are the most common problem, but other defects can depreciate the products and render them unsaleable, leading to significant economic losses for the producers and an increasing storage burden for the industry. The current system of bulk syrup classification is based on two distinctive parameters. One is based on physical properties such as light transmittance and soluble solids content (°Brix). The other is based on sensory appreciation by trained inspectors. How the various types of defects (Table 1) vary according to the collection period is unknown because the date of production is not systematically recorded. Sap composition variation can be in part attributed to metabolic changes occurring in deciduous trees during dormancy release or a stress response^16–18^.

**Table 1.**
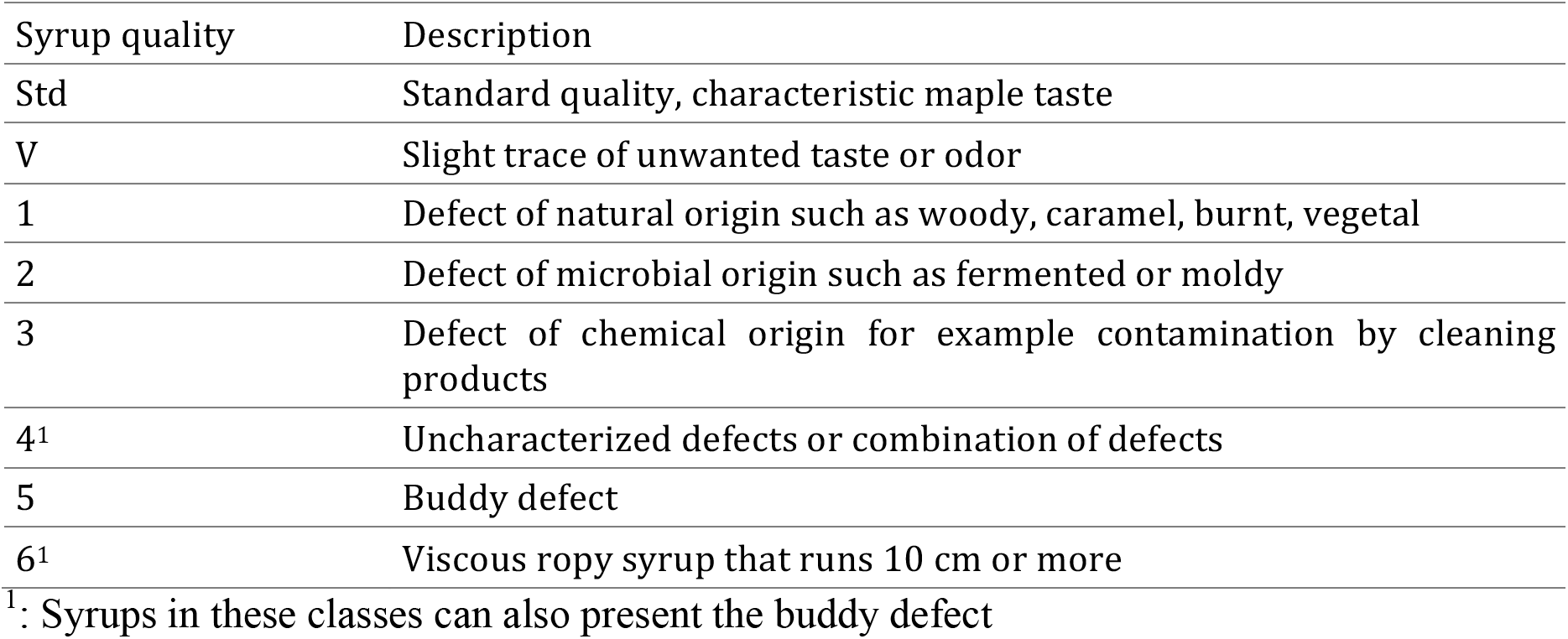
The official maple syrup classification system in Quebec.

The annual cycle of deciduous trees is composed of several steps including growth cessation, bud set, dormancy induction over the winter period, and its release during Spring. Specific metabolic changes have been reported during these steps^19–22^. Several molecules playing a role in dormancy are phytohormones such as gibberellin, ethylene, abscisic acid, cytokinins, and auxin^23–25^. Some of them are known to be transported to the plant xylem through the sap as biological signals to trigger metabolic changes during the annual cycle^23,24^. Some studies investigated dormancy release through untargeted approach such as transcriptomic profiling analyses, improving knowledge about the genetic network involved^26,27^. However, little is known about the metabolic changes occurring during dormancy release in maple trees and their relationship with sap properties^28–30^. Hence, access to commercially harvested maple sap from hundreds of trees and multiple locations during Spring offers a unique opportunity to study these changes. As the dormancy release period overlaps with the maple sap harvest period prior to bud break, molecules related to this metabolic change and that are transiting through the sap could directly or indirectly impact syrup quality. Moreover, the physiological requirement for sap flow and dormancy release may not be the same, such that these events are not necessarily synchronized. Sap flow has been reported to depend on temperature signals, cloud cover and rain^29^, while dormancy release and bud break depend on perception of photoperiodic and temperature signals^19,20^. In maple trees, a model was proposed by Raulier and Bernier^28^ based on winter chilling and spring warming records to predict the date of bud burst. This particular decoupling between sap flow and dormancy release is a conundrum for maple syrup production, because some years, the climatic conditions are such that sap flow is still heavy, but the tree physiology has reached a point where important quality defects appear in the syrup produced. In particular a characteristic “buddy” off-flavor, appearing around the time of bud burst, has traditionally marked the end of maple syrup production^30–33^, but its relationship to dormancy release has not been scientifically documented^34^. The fact that the prevalence of this specific defect appears to be unrelated to new technologies introduced over the last decades such as membrane processing, plastic tubing and vacuum utilization further supports the hypothesis that this defect is of biological origin^14,15^. Moreover, the potential to extend sap collection beyond this period is substantial and would benefit maple syrup producers.

Thus, we sought to identify the changes in maple sap in respect of dormancy release, associated with the appearance of quality defects, first with amino acid content profiling, then with a systems biology approach using a prototrophic yeast deletion collection as a tool for monitoring sap composition variation. We reasoned that the genomic tools available and the ecological association of *Saccharomyces cerevisiae* with deciduous trees makes them relevant and powerful biological reporters^7,35^. We adapted the approach by using the prototrophic yeast collection and a ureide-based control media to identify biological signals associated with sap composition, dormancy release and maple syrup quality. We found that changes in syrup quality coincide with the conversion of sulfate to organic sulfur in maple sap in the late stage of dormancy release.

## Results

### Relationship between sap quality and dormancy release

Our first aim was to improve our understanding of maple syrup defects with respect to the tree dormancy release. Our hypothesis was that a change in the tree metabolism is responsible for the buddy defect. Accordingly, we expected to see a relationship between a tree dormancy indicator and class 5 syrups, which corresponds specifically to the buddy defect (Table 1). We based our analysis on an index representing the remaining Sum of cumulative temperature necessary to reach the stage of Bud Burst (S_bb_), which we formulated using the model of Raulier and Bernier^28^, in which S_bb_ = 0 corresponds to bud burst. S_bb_ high value are therefore found early in the season, prior to bud burst. We expect this index to be more accurate for comparison across production sites and harvest years than the percent of cumulative flow. We related this index to the quality of samples using two approaches. First, considering only the percentage of class 5 syrups produced on the sampling day. Second, considering each defect classes independently.

In 2013, 66 sap samples were collected from nine production sites in the province of Quebec, Canada. The percentage of class 5 syrups was negatively correlated to the S_bb_ index (Spearman ρ −0.57, P-value= 7.7e^−7^). Over the Spring of 2016 and 2017, we obtained a total of 27 and 11 sap samples and 25 and nine syrups from 10 and eight locations, respectively (Supplementary data 1). For these samples, all types of defects classified by the industry (Table 1) were taken into account, by matching each sample with their syrup and submitting them to a standardized quality assessment procedure. Comparison between quality and S_bb_ index showed that standard syrups (Std) had significantly higher values than class 5 and class 6 syrups (Kruskal-Wallis P-value = 4.5e^−3^; Dunn test P-value = 0.01 (Std-class 5); 0.04 (Std-class 6), Supplementary Fig. 1 and data 1). No significant differences were observed among other defect classes. The results for both classification approaches are in agreement with our hypothesis and corroborate the previous observation that class 5 syrups appear shortly before the time of bud burst (class 5 S_bb_ range = 13-41).

### Amino acid profiling

Our second aim was to gather insight into maple sap composition variation over the dormancy release period to understand the origin of the buddy defect. We hypothesized that changes in amino acid composition are responsible for the defect appearance. Therefore, we quantified 36 amino acids or their derivatives in sap samples harvested in 2013 and searched for patterns that would associate their relative occurrence of the buddy defect (Supplementary Fig. 2, Supplementary data 1). The total content increased over the harvest period and was correlated with the S_bb_ index (Spearman ρ −0.52, P-value = 9.3e^−6^, Fig. 1a and1b). An individual correlation with the S_bb_ index was found for 23 amino acids or derivative out of the 36 quantified (Spearman, adjusted P-value >0.05) including 15 negative correlations (Supplementary data 1). The strongest correlations were found for valine, isoleucine, leucine and methionine (Spearman ρ −0.81 (adjusted P-value = 1e^−15^), −0.77 (adjusted P-value = 6e^−13^), −0.75 (adjusted P-value = 4e^−12^) and −0.69 (adjusted P-value = 2e^−9^, respectively)).

**Figure 1.**
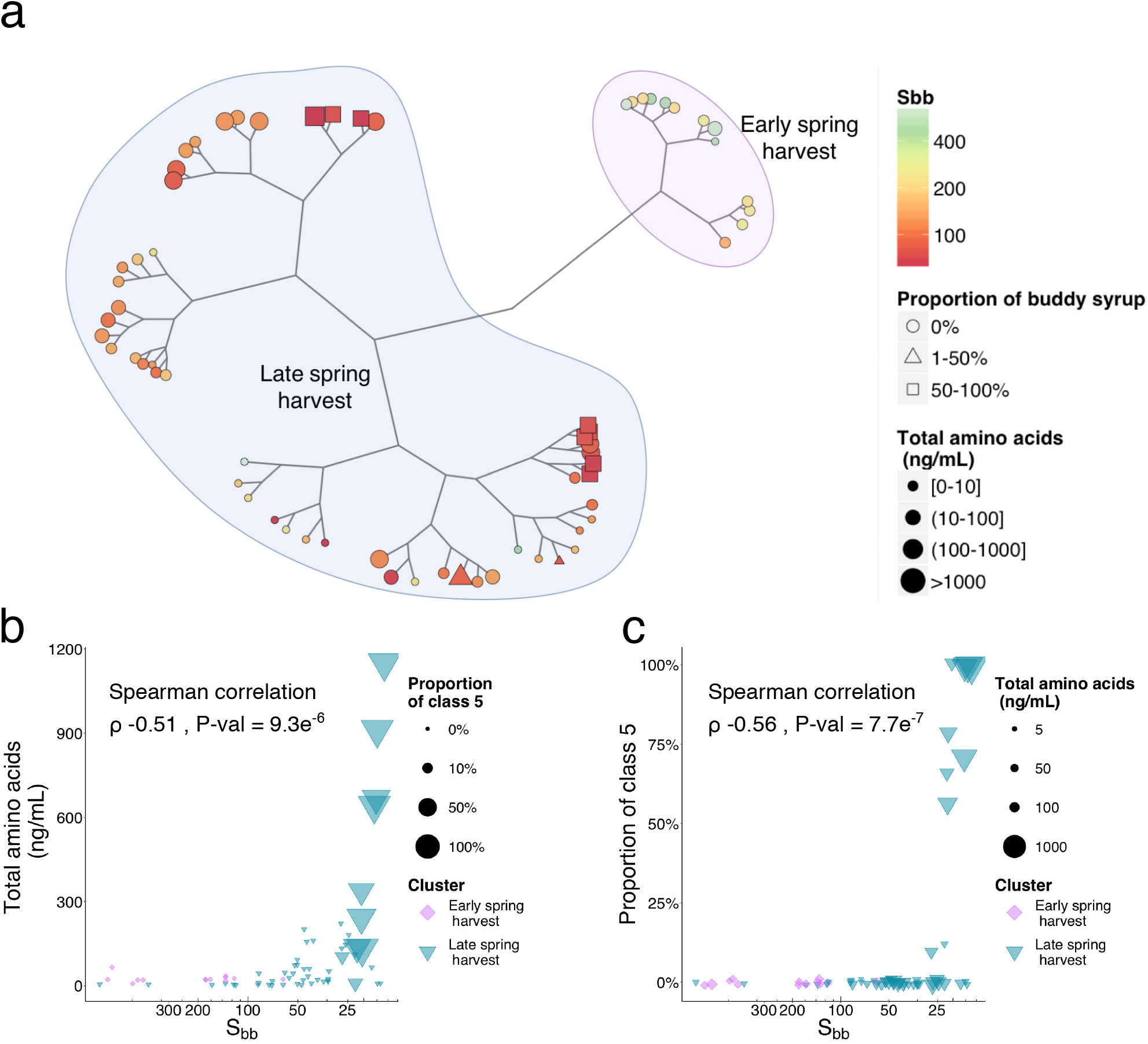
Maple sap amino acid profiles shift over the dormancy release process (a) Unrooted tree (Ward’ s method) of amino acid profiles of 2013 samples showing unsupervised clustering of early and late spring saps. Branch length represents square root distances. Color and shape size represent the S_bb_ index and total amino acid content, respectively. Shapes reflect the proportion of class 5 syrup produced on the day of sampling. (b) Class 5 syrup occurrence increases towards late Spring. Dot size reflects the total amino acid concentration. Colors and shape size refer to clusters defined in a) and total amino acid concentration, respectively. (c) Total amino acid contents increase towards late Spring. Dot size reflects class 5 syrup proportion.

The total amino acid and derivatives content was also correlated with class 5 occurrence (Spearman ρ 0.45, P-value = 1.4e^−4^). Correlations between the proportion of class 5 syrups and the 23 amino acids were also found (Spearman, adjusted P-value >0.05) including 14 positive correlations (Supplementary data 1). The strongest correlations were found for sarcosine, methionine, isoleucine and leucine (Spearman ρ 0.59 (adjusted P-value = 7e^−6^), 0.56 (adjusted P-value = 2e^−5^), 0.55 (adjusted P-value = 2e^−5^) and 0.54 (adjusted P-value = 2e^−5^), respectively). Given that sarcosine and methionine are part of the plant one-carbon metabolism,^36^ these results point to a link between this biological process and the buddy defect.

Since amino acids can compete as a substrate in Maillard reactions that occur during the transformation process of maple syrup and contribute to its flavors, their relative proportion was also considered. A hierarchical classification of amino acid proportion profiles shows that samples clustered according to the S_bb_ index (Wilcoxon P-value = 9.4e^−7^), but not by production sites (Fig. 1a, Supplementary data 1). According to spring harvest clustering sample, the amino acid profiles show a breaking point at around 100 S_bb_, which is before the appearance of class 5 syrups (Fig. 1c). The class 5 associated samples were further split in two groups within the late spring harvest cluster, indicating that at least two subtypes of amino acid profiles can lead to the buddy defect, highlighting the complexity of this matrix.

### Yeast fitness competition in sap

To further our aim to gather insight into maple sap composition variation over the dormancy release period and its relationship with maple syrup quality, we used the prototrophic version of the barcoded yeast deletion collection as a biological reporter. The collection represents a set of genome-wide non-essential individual gene deletions. Because the yeast genome is well annotated, an enhanced or diminution of growth defect for a particular strain or group of strains can indicate which molecules are present in the media. We thus captured the variation in yeast growth in 27 saps from 2016 in the form of a yeast genome-scale fitness profile of 4090 non-essential genes (Supplementary data 2). We compared strain fitness in a minimal control media containing sucrose and allantoin as carbon and nitrogen source, respectively, to fitness in media in which the sucrose was substituted with the sap samples. Since we have previously shown that allantoate, an intermediate in allantoin catabolism, is the main nitrogen source for yeast growth in maple sap during the spring flow period^9^ and sucrose is the main carbon source, we expected to emphasize the effect of other relevant compounds under these conditions. A principal component analysis (PCA) on correlations between the fitness profiles showed differences between sap samples and the sucrose controls and that standard (Std) quality sap can also be discriminated from buddy sap (class 5) with this approach (Supplementary Fig. 3). Hierarchical classification of fitness profiles shows that sap samples clustered according to the S_bb_ index (Wilcoxon P-value = 3.6E^−4^) (Fig. 2a) with a breaking point around 30-40 S_bb_ which coincides with the appearance of various defects, class 5 in particular (Fig. 2b).

**Figure 2.**
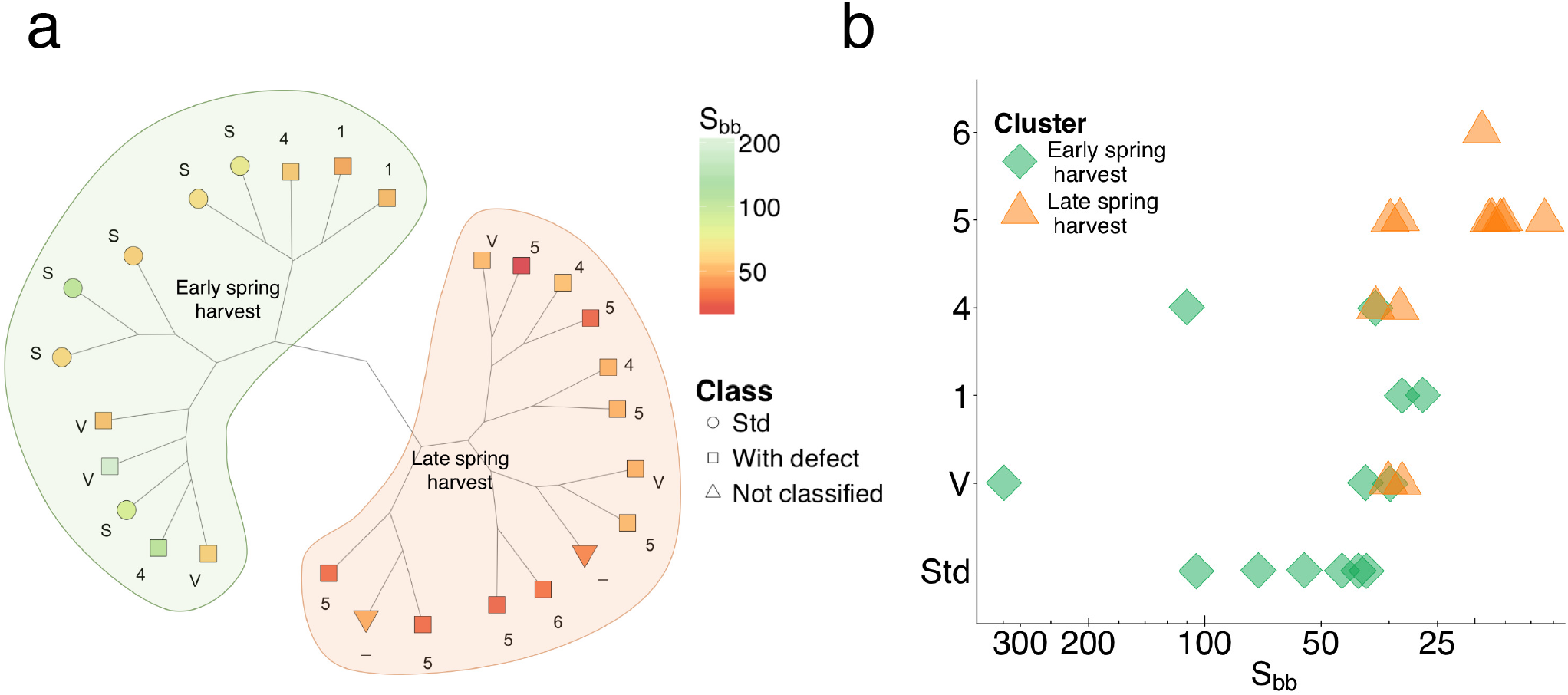
Yeast fitness profiles shift according to dormancy release and maple syrup quality (a) Unrooted tree (Ward’ s method) of the yeast fitness collection profiles of sap samples from 2016 showing a partition between Std and class 5 samples. Branch length represents square root distances while color and shape represent S_bb_ and sample quality, respectively. (b) Sap defects are not randomly distributed over the dormancy release period. Buddy defects appear after a turning point coinciding with the clustering based on yeast fitness profiles. Color and shape represent clusters defined in a).

Based on the fitness profiles, we selected three lists of deletion strains: (1) the sap list, reports strains whose fitness distribution significantly differed between the control media and the sap (Kolmogorov–Smirnov test, adjusted P-value < 0.05, Supplementary data 2), (2) the S_bb_ list contains strains whose fitness is correlated to S_bb_ (Spearman test, adjusted P-value < 0.05) and (3) the quality list, for which strain fitness differed between the Std and the class 5 samples (adjusted P-value < 0.05 & Δfitness ± 0.02). The sap, S_bb_ and quality lists contained 1662, 1035 and 218 strains, respectively (Supplementary Fig. 4), including 204 strains common to the S_bb_ and quality lists. The three lists were sharing two Gene Ontology Biological Process (GO-BP) enrichments, electron transport and mitochondria translation. We found more common categories for the quality and the S_bb_ list such as amino acid activation, RNA processing, mitochondrial activity and sulfate assimilation (Permute score < 0.05) (Fig. 3, Supplementary data 3). Gene ontology enrichments were compared with results from VanderSluis *et al*., (2014)^37^ that report the response of the same *Saccharomyces cerevisiae* prototrophic deletion collection to sole carbon or nitrogen sources. Since our competition media was supplemented with allantoin to ensure sufficient growth in sap and we observed that other nitrogen sources as amino acids become available towards bud break (Fig. 1b), we expected biological processes involved in allantoin utilization to be enriched in our lists. This was indeed the case, 14 enrichments were common between the results of VanderSluis and our lists (Supplementary data 3). Aside from these common enrichments, the sap list shares most of their enrichments with genes involved in the utilization of glutamate (Fig. 3), the most abundant amino acid in maple sap (Supplementary Fig. 2). The quality and S_bb_ lists share most of their enrichments with substrates that require mitochondrial functions for their utilization, namely ribose, galactose and proline^38,39^. The enrichment for sulfate transport appears to be unique to the S_bb_ index and sulfate assimilation to both the S_bb_ and quality list. An analysis of gene-metabolite association showed that our three lists were enriched for genes associated with specific metabolites (Supplementary data 3). In particular the S_bb_ was associated with sulfate, while the quality list was associated with sulfite, phosphoadenosine phosphosulfate (PAPS) and S-adenosylmethionine (SAM) (Fig. 4, Supplementary Fig. 5). Altogether these results indicate that changes in sulfur composition in maple sap is associated with dormancy release and maple syrup quality.

**Figure 3.**
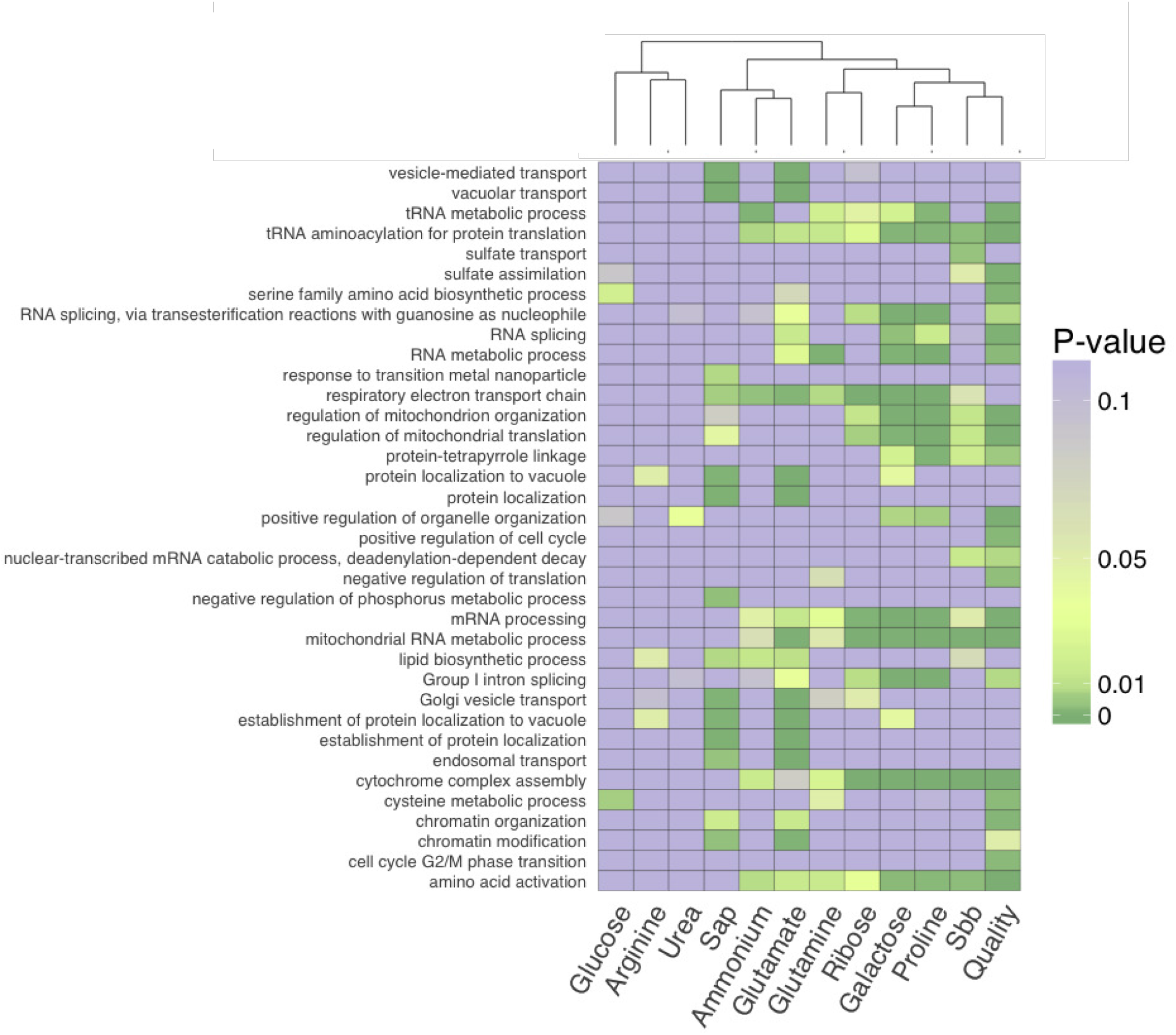
Heatmap of the most enriched biological processes in Maple sap, Quality and S_bb_ lists (P-value < 0.01) alongside enrichments for genes involved in utilization of particular substrates^37^. We excluded the enrichments related to growth on allantoin common between our assay and VanderSluis experiments (P-value < 0.01).

**Figure 4.**
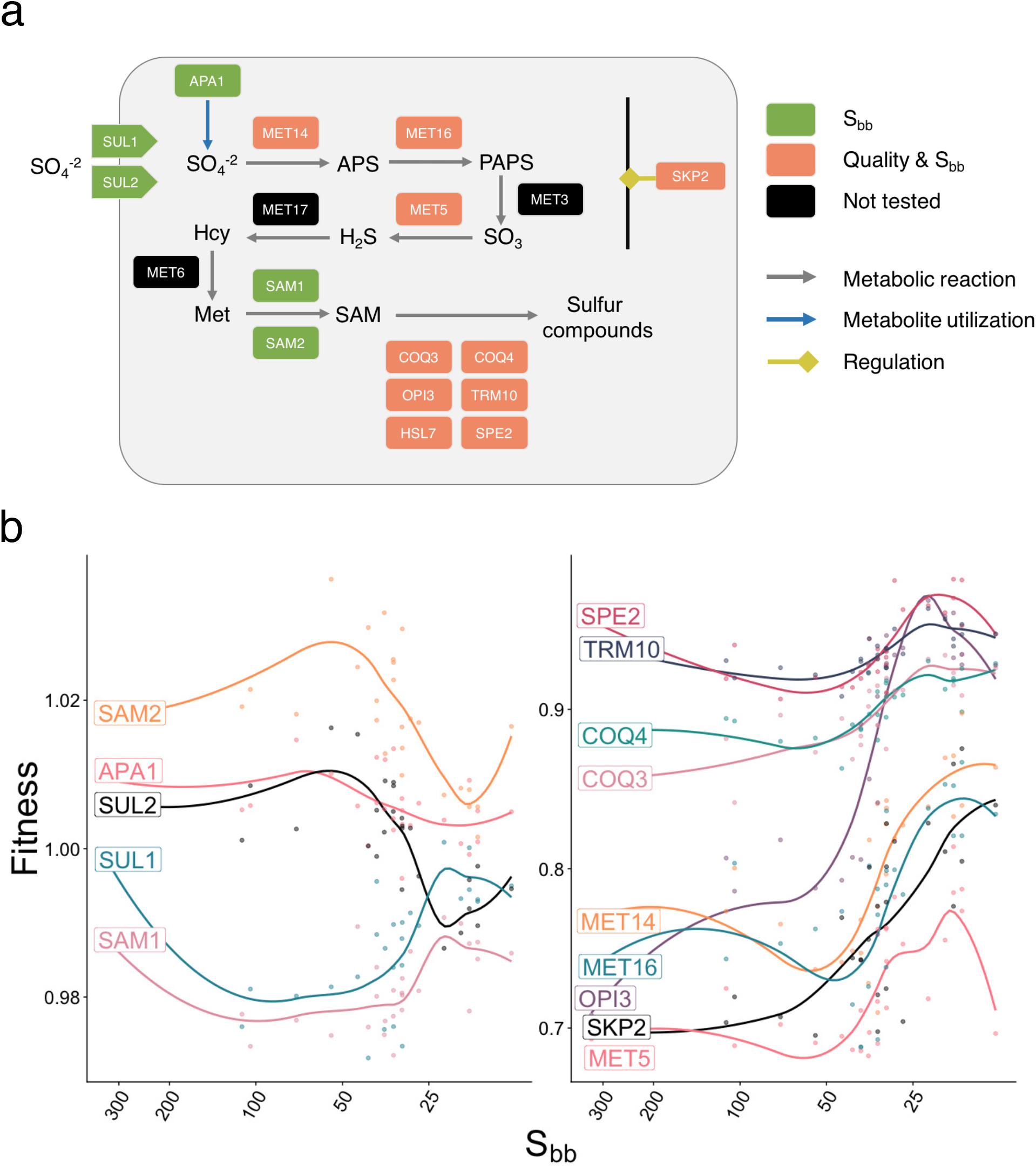
The *S. cerevisiae* deletion strains involved in the sulfate metabolic pathway and its regulation are enriched in the S_bb_ and quality lists. (a) Schematic representation of the sulfate metabolic pathway. APS: Adenosine phosphosulfate; PAPS: Phosphoadenosine phosphosulfate; SAM; S-adenosyl-L-methionine; Met: Methionine; Hcy: Homocystein. (b) The detailed fitness effect of each gene deletion in relation to the S_bb_ index of maple samples. SUL1 and SUL2 gene deletion strains present opposite fitness as they are low- and high-affinity sulfate transporter, respectively. Line represent the loess regression curve between the fitness and the S_bb_ index.

To confirm the results of the yeast fitness competition, we performed individual liquid growth assay for four strains (*opi3, met14, met5* and *skp2*) involved in the sulfur metabolism pathway (Fig. 4, Supplementary Fig. 4) in the 2016 and additional 2017 samples. The *opi3* strain growth profile matched that of the fitness competition experiment for the 2016 samples, but growth in additional samples from 2017 did not support the relationship with quality (Supplementary data 4). No clear pattern was found in *met14* regarding the S_bb_ value or the quality indicating that interaction with other strains may have been key to the results obtained in the fitness competition experiment (Fig. 5a). However, the strains *met5* and *skp2* had minimal or no growth in all Std saps but were able to grow in all class 5 saps (Fig. 5a). Since the *met5* deletion strain requires organic sulfur for growth, variation in its content in maple sap may explain the appearance of the buddy defect.

**Figure 5.**
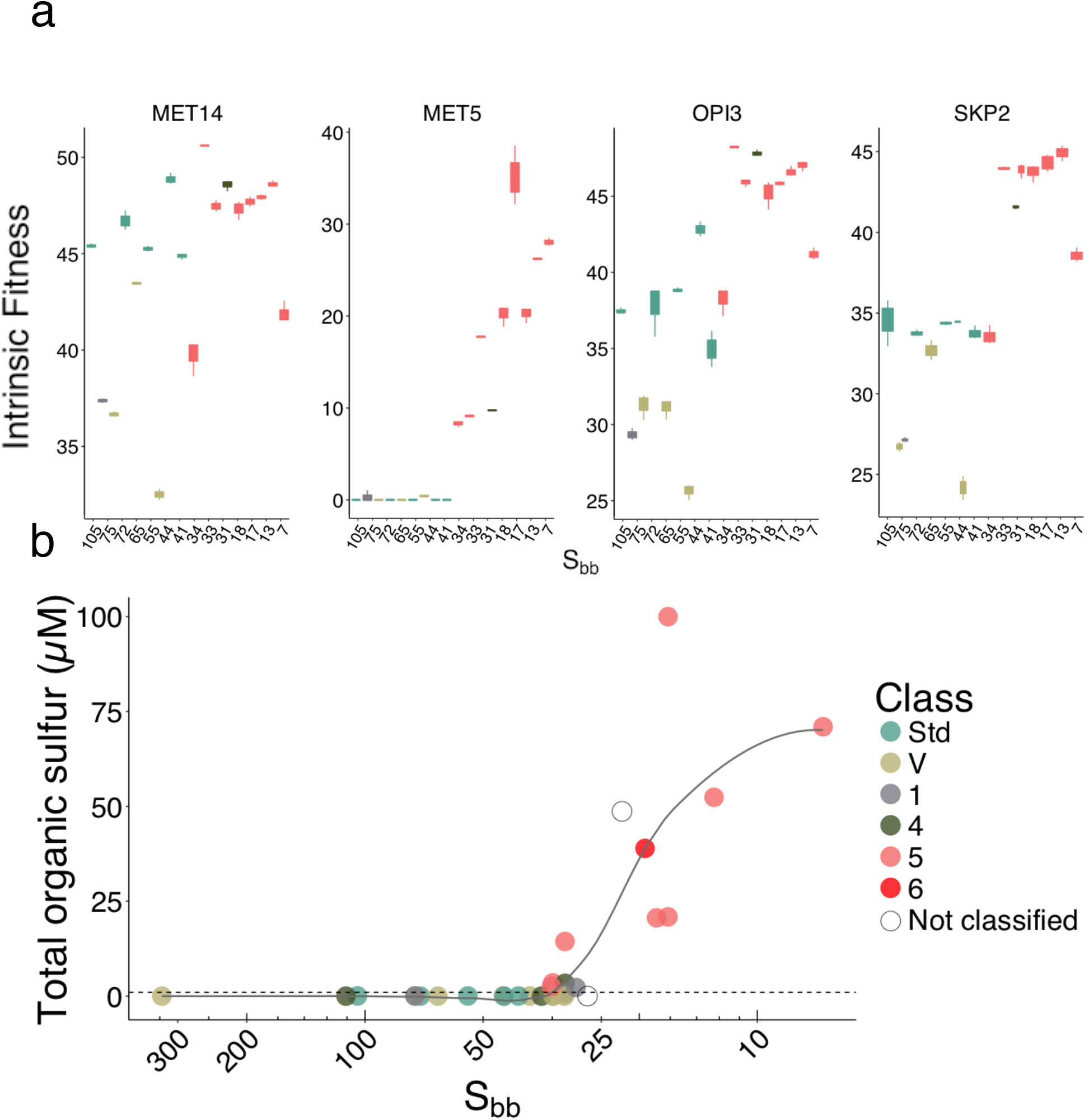
Growth profile of *Saccharomyces cerevisiae* strains impaired in their sulfur metabolism pathway in a subset of 2016 and 2017 samples. (a) Intrinsic fitness of the four mutant strains *met14, met5, opi3* and *skp2* in 2016 and 2017 saps. Colors represent quality classes. X axis represent samples ordering by decreasing S_bb_. (b) Organic sulfur concentration in saps, as estimated by a standard curve of *met5* intrinsic fitness in methionine, varies along the sample S_bb_ and coincides with quality. Line represent the loess regression curve between the estimated organic sulfur concentration and the S_bb_ index.

We used the *met*5 strain growth as a proxy of estimate the organic sulfur content based on a methionine standard curve (Fig. 5b). We found a significant correlation between the estimated organic sulfur concentration and the S_bb_ index (Spearman ρ −0.70, P-value: 1.2e^−05^, Fig. 5b) and with the sap quality (Kruskal-Wallis P-value: 2.8e^−3^; Dunn test P-value = 0.02 (Std-class 5)), reflecting a change in the nutrient content of maple sap associated with dormancy release. Moreover, the organic sulfur increase observed between 30 and 40 S_bb_ coincides with the occurrence of class 5 syrups in 2016 and 2017. Thus, at least for these samples, *met5* appears to be an appropriate biomarker of maple sap quality with an interval concentration of 2.5μM to >100μM.

### Sulfur compounds in maple sap

The yeast fitness competition revealed a specific association between dormancy release and sulfate, while the buddy defect was associated with sulfate assimilation products. Therefore, we hypothesized that the maple tree metabolism is releasing sulfate that can be metabolized by the sap microbiota in a variety of organic sulfur molecules, including amino acids such as methionine. We measured inorganic sulfur (SO_4_^2−^), Ammonium (NH_4_^+^), and phosphorus (PO_4_^3−^) in saps harvested in 2016 and 2017. Ammonium was measured because it is a catabolic product of allantoate and would therefore reflect microbial activity, and phosphorus was measured as a negative control, since the negative regulation of phosphorus metabolic process was specifically enriched in the sap list, but not in the S_bb_ or quality list, SO_4_^2−^ and NH_4_^+^ were indeed correlated wih each other (ρ 0.78, P-value: 2.6e^−6^) and significantly higher in class 5 samples than in Std samples (SO_4_^2−^: Kruskal-Wallis P-value = 7e^−4^; Dunn test P-value = 0.05 (Std-class 5); NH_4_^+^: Kruskal-Wallis P-value = 5e^−3^; Dunn test P-value = 4.8e^−4^ (Std-class 5); 6.7e^−3^ (Std-class 6); Fig. 6a; Supplementary data 4). We also found a correlation between NH_4_^+^ and the organic sulfur content (ρ 0.61, P-value: 9.6e^−4^). Their concentrations were also highly correlated with the S_bb_ index (Spearman ρ −0.80, 8.9e^−7^; ρ 0.86, P-val =1.5e^−8^, respectively) (Fig. 6b) with the correlation between SO_4_^2−^ and the S_bb_ index being the strongest reported in this study. As expected, the PO_4_^3−^ concentration was not correlated to the S_bb_ index or the quality (Fig. 6). Altogether, these results support the conclusion that the organic sulfur content in maple sap results from microbial activity.

**Figure 6.**
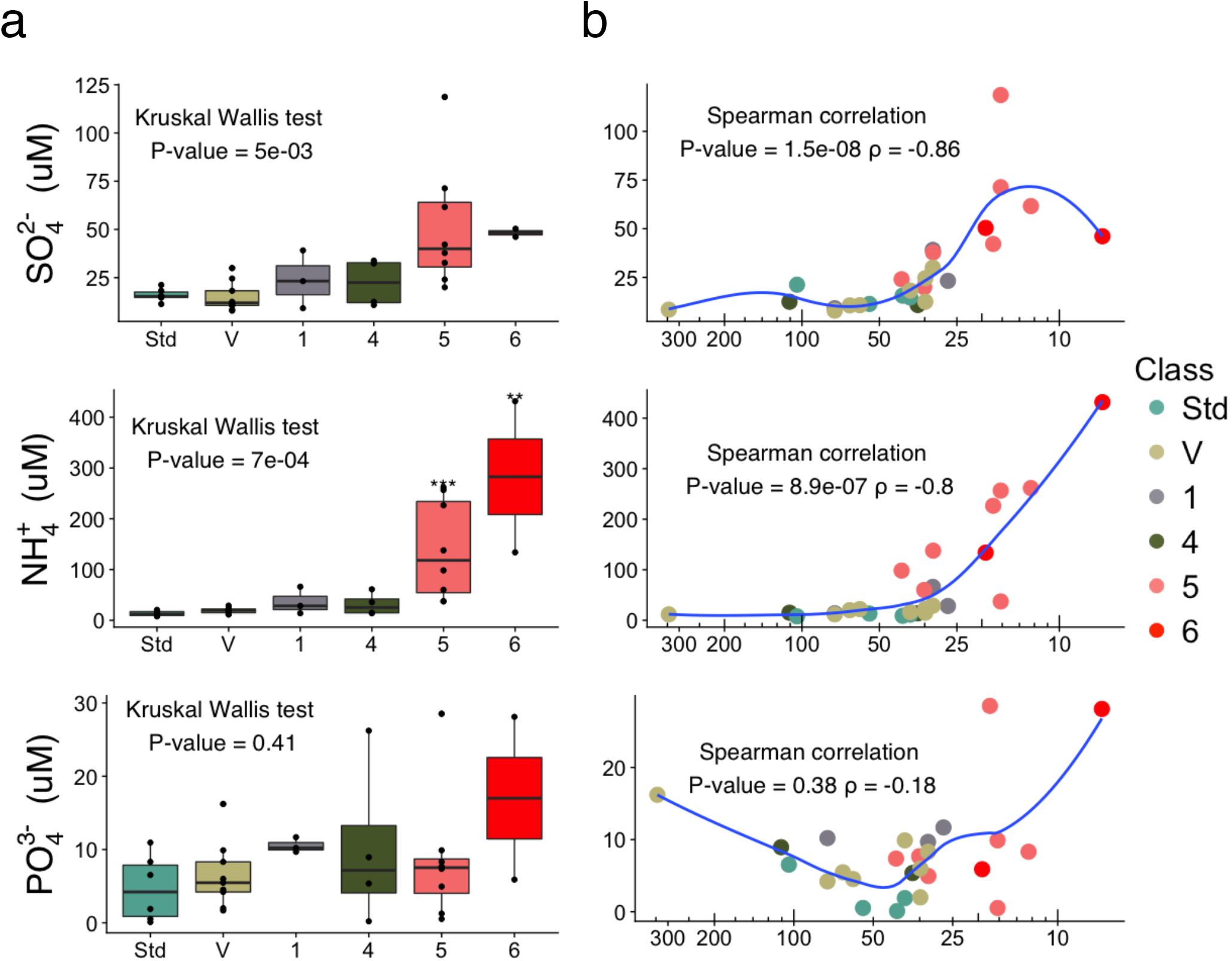
Variation in NH_4_^+^, SO_4_^2−^, and PO_4_^3−^ concentration in maple sap (a) Boxplots of concentration by sample quality as defined in table 1 showing that NH_4_^+^ and SO_4_^2−^ are significantly different between Std and class 5 syrups. Defect syrups were compared to the Std group by a Dunn’s test after a Kruskal Wallis test (P-value < 0.05). *denotes a significant difference (P-value < 0.05). Boxplots show the median, 1^st^ and 3^rd^ quantiles. (b) Sap variation in NH_4_^+^ and SO_4_^2−^ are correlated to dormancy release (S_bb_) (Spearman P-value < 0.05) while PO_4_^3−^ is not.

As the concentration of organic sulfur in sap was discriminant between Std and class 5 samples, and that the range of concentration observed for sulfur amino acids in 2013 (methionine, cysteine (2 sulfur atoms), cystathionine and thiaproline) was lower (range 20nM to 28.5μM, Supplementary data 1) than the estimated range of total organic sulfur, we looked for additional organic sulfur compounds in maple sap. We targeted three molecules as likely candidates, SAM, S-Adenosyl Homocystein (SAH), and Methylthioadenosine (MTA). We based this choice on the enrichment results (Supplementary data 3), and the fact that in plants, SAM can be converted to MTA to produce ethylene, which has been shown to have a concentration that peaks in late Spring in a maple tree species^40^. We quantified SAM, SAH and MTA, but the three molecule concentrations were below the limit of detection in all sap samples regardless of their quality (detection limits SAH & MTA: 2.6 - 2.9nM; SAM: 33nM; Supplementary data 4). We also measured SAM concentration by generating the double yeast mutant strain *sam1 sam2* as this strain is auxotroph for this molecule. The auxotroph strain did not grow in any of the samples, suggesting the absence or very low concentration of SAM (<15nM) (Supplementary data 4).

## Discussion

Maple syrup is a complex natural product whose attributes result from the biological, microbial, and human activity, which lead to variation in quality and functional properties that in turn can have drastic consequences for marketability^3,6,12,31–33,41–43^. However, little is known about the exact molecular compounds that explain this quality variation, and even less about the precursor compounds present in the sap, as most of the defects can only be detected once the syrup is produced^34^. The collection of sap and the transformation process are not standardized, making it challenging to identify factors impacting the final quality of the product. On the one hand, the classification currently in use is not well suited to investigate the origin of specific defects since some of them can be found in multiple classes. For instance, the buddy defect can be classified in class 4, 5 or 6 according to its intensity or combination of defects. One the other hand, the mechanisms of dormancy release and bud burst in deciduous trees are beginning to come to light^19,20,22,26,27,44^. Here we devised a basic dormancy index to compare our samples. So far, the index only takes into account climatic parameters from the nearest meteorological station, therefore there is much room for improvements with factors associated with dormancy release, such as considering the effect of photoperiod, altitude, age and genetic populations of trees^19^. Despite these pitfalls, we showed a consistent relationship between the tree dormancy release and the defect appearance, as it recurrently appears at S_bb_<41, supporting earlier empirical observations^34^. Our model should prove more accurate than the harvest date to predict the appearance of the buddy defect regardless of the harvest year. Nevertheless, curative methods need to be developed to prevent the premature ending of the harvest season, therefore understanding the changes leading to the appearance of the defect is necessary.

Our first efforts to identify the underlying cause of the buddy defect were to profile 36 amino acids in maple sap samples collected over the Spring. Amino acids are precursors for the Maillard reaction, a non-enzymatic browning reaction that occurs during the transformation process of syrup and contributes to the development of the flavors and colors of the product^41^. Our results confirmed a global increase of amino acids content in sap prior to bud burst, which has been reported in pioneer work^9^. We found significant correlations between the S_bb_ index and amino acid contents, but correlations were overall weaker with class 5 syrup occurrence. Since we only obtained information for class 5 syrup produced on the sampling day and not matching samples in 2013, the possibility remains that a more significant effect was missed. Considering that there could be competitive effects between amino acids as the substrate in the Maillard reaction,^45^ we also considered their relative proportion profiles. The results show a main clustering split at around S_bb_=100. These results indicate a first shift in amino acid profiles unrelated to the appearance of class 5 samples. Interestingly, a shift in microbial communities between early and mid-season has previously been reported^1,8^ and could be related to the pattern observed here. The class 5 samples were also grouped in two sub-clusters within the low S_bb_ cluster, indicating at least two subtypes of amino acid profiles can be associated with the buddy defect. Hence, it is likely that class 5 syrups represent two types of defects both recognized as buddy or that the conditions leading to the same defect development in syrup are not unique.

To further understand changes in maple sap composition related to dormancy release and syrup quality, we used an untargeted approach using the yeast *S. cerevisiae*. Yeast fitness profiles clustered similarly to amino acid and their derivate profiles, partitioning early and late harvest samples. These results show the usefulness of this approach to identify compounds that vary in coordination with dormancy release. In addition, the S_bb_ breaking point matched a shift in syrup quality, indicating that this approach revealed a different signal than amino acid profiling. When comparing biological process enrichments between our lists of responsive strains and lists of strains with altered growth rate on various substrates^37^, we found that we could explain most enrichments. However, enrichment for sulfate transport and sulfate assimilation were specific to our lists, hinting at the release of sulfate over dormancy release and its assimilation into organic compounds in buddy saps. Indeed, sulfur appears to be necessary for the bud break as various sources of sulfur compounds are increasing prior to this event, depending on the tree species^46–48^. Several sulfur compounds (cysteine, glutathione, S-methyl-methionine, sulfate) transiting through the xylem prior to bud breaks have been reported in woody plants, however composition varies between species^46–49^. We confirmed the presence of sulfate in maple sap and showed an increase in sulfate towards bud break. Also, based on the growth of *met5*, a strain auxotroph for organic sulfur, we showed that organic sulfur is not available in Std saps and its appearance coincides with that of the buddy defect. For SO_4_^2−^ to be incorporated into organic compounds, a nitrogen source would also have to be available. Given that allantoate was shown to be the major source of nitrogen available to yeast in maple sap and increased in concentration over the harvest period, and that microbial contamination also increases^2,6^, we measured the allantoate degradation product NH_4_^+^ to assess microbial activity. We found that NH_4_^+^ and organic sulfur are correlated and associated with quality, which most likely indicates that microbial activity is responsible for the presence of organic sulfur and suggests that microbial activity plays a role in the buddy defect appearance.

These results, combined with the knowledge that the buddy defect cannot be detected before boiling the sap^34^, suggest that one or a combination of organic sulfur compounds produced by microorganisms are precursors to the reaction generating the buddy off-flavor during the transformation process. Sulfur compounds are known to be associated with negative flavors in various food^50–52^. The main sulfur amino acid measured in our assays was methionine and was also one of the most correlated to class 5 syrups. However, its concentration range was lower than the estimated organic sulfur content, suggesting the presence of additional molecules. Therefore, we looked for other sulfur compounds enriched in our results, namely SAM and SAH, and the related metabolite MTA. SAM is the precursor of ethylene, and MTA is a by-product of the reaction^53^. Ethylene is a phytohormone proposed to be involved in dormancy release and bud breakthrough crosstalk with abscisic acid^19,54^. In *Acer platanoides* L, a seasonal increase in ethylene coinciding with bud burst has been observed^40^. However, we did not detect the presence of these metabolites in maple sap. Given the central metabolic role of SAM in sulfur metabolism^55^, it is possible that the effects observed on yeast fitness are the result of a convergence of the metabolism of multiple organic sulfur sources. Furthermore, as it is collected, maple sap nutrient becomes available to microorganisms, combining the molecules produced by the tree and others resulting from microbial activity. It is reasonable to suspect that methionine, the main organic sulfur source in maple sap, is produced by microbial activity. Prior studies on maple sap microbial communities over the flow period did not report on buddy syrups nor nitrogen sources. Therefore, additional information should be gathered to attempt to capture the microbial communities and their functions involving sulfur compounds. Interestingly, a method to prevent the production of buddy syrups has been patented in 1963 and involved incubation with an inoculum of *Pseudomonas geniculata*^3^. However, the biochemical mechanism of action was not reported and the method was not implemented in the industry.

Overall, our approach enabled us to contribute towards the existing knowledge of sap composition variation in relation to dormancy release and maple syrup quality. We devised a dormancy index that could be a useful tool to predict syrup quality and we report that the organic sulfur content, NH_4_^+^ and SO_4_^2−^ are potential markers to monitor sap quality. We conclude that while the dormancy release certainly contributes to a coordinated appearance of the buddy defect, we suspect a determining role for microbial activity. Are there specific microbial communities or community functions associated with the buddy defect? The interactions between microbial communities, environmental conditions and sap composition are promising avenue of research to further understand maple syrup quality variation and develop biocontrol applications.

## Methods

### Maple sap and syrup samples

Maple sap concentrated by membrane processing and syrups were obtained from Québec producers during the Spring of 2013, 2016 and 2017 with an emphasis on the end of the flow period. Samples were kept frozen at −20°C until analysis.

One sap sample was obtained on each processing day during the entire 2013 season from nine production sites. The concentration of solid content for each sap sample was measured with a digital refractometer (0-95 solids or °Brix, Reichert AR200). All barrels of maple syrup produced by nine production sites were subjected to sensory evaluation by certified inspectors following standard procedures used in the maple industry. The percentage of class 5 syrup barrels, corresponding specifically to the buddy defect, produced on each sampling day was calculated. For sample harvested in 2016 and 2017, sap concentrates were obtained with the corresponding syrup produced. Sap concentrates were centrifuged for 5 min × 915 g at 4 °C and sterilized by 0.2 μM filtration. The concentration of each filtered sap was measured with a digital refractometer (°Brix). Syrups were subjected to sensorial evaluation by three certified inspectors and classified as of standard quality (Std) or otherwise with the defect types presented in Table 1.

### Index of dormancy release

Meteorological records from the nearest stations were obtained from the Canadian government public records (http://climate.weather.gc.ca). Missing data were imputed using data from the nearest station. The observed temperature sum (S_w_) was calculated by cumulating from December 1^st^ the positive difference between the average daily temperature and 10°C. Similarly, the number of chilling days (d_C_) was obtained by cumulating the number of days on which the average temperature was below 10°C. A combined chilling and warming model^28^ was used to predict the temperature sum necessary for leaf emergence (S_wp_) given the d_C_ observed for each sample. In this model, leaf emergence corresponds to a bud burst state where leaf margins are visible but not yet unfolded^29^. As an index of dormancy release at the time of sampling, we used the difference between S_wp_ and S_w_ to estimate the remaining temperature sum necessary for this stage of bud burst (S_bb_). In our data, S_bb_ is highly correlated with the Julian date (Spearman ρ −0.99, P-val = 2.2e^−16^).

### Culture media

The control media for our experiments consisted of 1.75 g.L^−1^ of yeast nitrogen base, 1.25 g.L^−1^ of allantoin and 2% sucrose. For the sap media, sucrose was substituted by sap concentrates diluted to a final density of 2°Brix.

### Functional genomic screen

The yeast deletion collection used was obtained from VanderSluis et al. 2014^37^. The functional genomic screen was performed as in Filteau et al. 2017^7^, with the following modifications: liquid assays were carried out in 250 μL of control and sap media in 96 well plates, after an average of 18 generations, aliquots from eight replicate wells were pooled for DNA extractions. A total of 50 libraries, divided into two runs, were then constructed by PCR, including four from the initial strain pool, 18 sucrose controls and 27 saps from 2016, using the primers listed in Supplementary data 1. DNA extractions, PCR, library preparations and sequencing were performed as in Filteau et al. 2017^7^.

### Sequencing analysis

Sequencing results were analyzed as in Filteau et al. 2017^7^, at the Plateforme d’Analyses Génomiques of the Université Laval (IBIS), with the following modifications: the sequence reads were mapped to the reference using Geneious R6^56^, with the following parameters: trim including region from 1 to 80 first nucleotides, word length 20, index word length 15, maximum mismatches per reads: 4%, allow gap 1%, maximum gap size 1 and maximum ambiguity 16. Fitness calculation was normalized using the following pseudogenes strains: “YLL017W”, “YIL170W”, “YCL075”, “YFL056C”, “YIL167W”, “YIR043C”. After alignments, library with more than one million reads were used for further analysis, excluding one initial pool and three sucrose controls. We obtained an unambiguous assignment for 83% of reads and Spearman correlation coefficients of ρ 0.92 to 0.93 between initial pool replicates and ρ 0.92 to 0.97 between sucrose replicates.

There were 4772 strains present in the prepared pool, among which 4484 were detected. The 4090 strains that had a sum of more than 100 reads in the three initials pools replicates were considered for further analysis.

A principal component analysis (PCA) was performed on the strains fitness according to the sap quality using the R packages FactoMineR and factoextra. Two complementary statistical analyses were then performed on strain fitness. We first evaluated the correlation between the S_bb_ index and the strain fitness using a Spearman correlation test and a Welch’ s test on the five Std saps and six buddy saps. Gene Ontology (GO) and metabolites enrichment of each list, including VanderSluis dataset^37^, on the *S. cerevisiae* prototrophic collection was performed using GoElite with the same parameters as in Filteau et al. 2015 retrieved form Saccharomyces Genome Database and Yeast Metabonome DataBase, respectively^57,58^.

### Growth experiments

We selected four mutant strains (*met5, met14, skp2* and *opi3*) from the prototrophic collection to validate the functional genomic screen results. Each strain was individually grown overnight at 30°C without agitation in the control media, then, they were inoculated in fresh sap media. Initial Optical Density (OD) was adjusted for each strain at 0.05 in 200 μL in a 96 well microplate and measured in triplicates. OD was measured each 30 minutes for 36 hours at 30°C using a Tecan plate reader (Zürich, Switzerland). For *met5* strain, after modelling the growth curve for each well using Gompertz model fit in JMP13 software (SAS Institute, Cary, NC), the intrinsic fitness value was calculated by integrating the area under the curve up to 48 hours of growth and used to estimate the total organic sulfur based on a standard curve of methionine (1mM to 10nM).

### Strain constructions

To generate the double yeast mutant strain *sam1/sam2*, the SAM2 locus was replaced with the HPH antibiotic resistance cassette on *sam1* competent cells strain. The deletion cassette was designed and amplified with the plasmid pFA6-hphNT1^59^, and the *SAM2* flacking region with the following oligo sequences (supplementary data 4). Cells regeneration was done on selected media YPD supplemented with the antibiotics 250 μg.mL^−1^ of hygromycin, 200 μg.mL^−1^ kanamycin and 60μM of SAM (Sigma Aldrich A7007) since the double mutant strains is auxotroph for this molecule. To verify the cassette insertion, amplification was performed using the Forward primer and an internal cassette primer (Supplementary data 4).

### SAM, SAH and MTA detection

Concentrated saps for harvested 2016 and 2017 year were filtrated by a micropore of 0.2 μm before quantification. SAH, SAM were purchased from Toronto research chemicals (Ontario, Canada) and MTA form MTA Cayman chemicals (Ann Arbor, MI). Acetonitrile LC-MS and acetic acid LC-MS of UPLC-MSMS grade were purchased from VWR International (Quebec, QC, Canada) and Fisher Scientific Ltd (Montreal, Quebec), respectively. UPLC-MSMS analysis was performed using a Waters Acquity H-Class Ultra-Performance LC system (Waters, Milford, MA, USA), equipped with a quaternary pump system. Samples were loaded on an Aquity BEH Column C18, 1.7 μm, 2.1 mm × 50 mm from Waters set at 30°C. The mobile phase consisted of 100% aqueous acetic acid (glacial) pH 2.6 (eluent A) and acetonitrile 100% (eluent B). The flow rate was 0.35 ml.min^−1^ and the gradient elution was 0 −1 min, from 0% to 0% B; 1-3.5 min, from 0% to 100% B; 3.5-4 min, from 100% to 100% B; 4-6 min, from 100% to 0% B.

The MS analyses were carried out on a Xevo TQD mass spectrometer (Waters) equipped with a Z-spray electrospray interface. The analysis was performed in positive mode. The ionization source parameters were capillary voltage 3.0 kV; source temperature 120°C; cone gas flow rate 0L.h^−1^ and desolvation gas flow rate 500 L^−1^; desolvation temperature 250°C. Multiple reaction monitoring (MRM) transitions, individual cone voltages and collision energy voltages were summarized in Supplementary data 4.

Data acquisition was carried out with MassLynx 4.1 (Waters). Quantification was performed based on external calibration.

### Statistical analysis

Statistical analysis was performed using R for Spearman correlation, Kolmogorov–Smirnov, Welsh, Anova, Tukey, Kruskal-Wallis and Dunn test. Significant stains form the Spearman correlation S_bb_ list and Kolmogorov–Smirnov test sap list were adjusted P-value < 0.05. For the quality list, significant strains from the Welsh test was adjusted P-value < 0.05 and Δfitness ± 0.02.

### Amino acid quantification

Amino acids analysis was conducted on LC/MS/MS system according to EZ:faast protocol for free amino acids (Phenomenex, Torrance, CA, USA). The EZ:faast kit contains standard solutions, reagents, sorbents tips and LC column. Chemicals and additive such as methanol (LC-grade) and ammonium formate (Mass apectrometry grade) were purchased respectively from Fischer Scientific and Sigma Aldrich. Water used for analysis was obtained from MILLI-Q Water System. For samples analysis, 200μL of maple sap was used for SPE extraction and derivatisation of amino acids following EZ:fasst protocol. Then, after solvent evaporation, the extract was reconstituted in 100μL of the initial LC mobile phase, and 2μL of this sample was injected on LC-MS/MS system. The LC system (Prominance, Shimadzu) is equipped with binary pumps LC-20AB, degasser DGU-20A5, column oven CTO-20 AC and an Autosampler SIL-2. The Qtrap3200 Mass Spectrometer Detector (Applied Biosystems, Sciex) was used for MS/MS analysis. The MS parameters are provided by Phenomenex (EZ:faast kit) and have been used without significant modifications. Amino acids content was then analyzed according to its concentration (μM) and relative mass proportion (mg.L^−1^) in each sample harvested in 2013 adjusted to 2°Brix. Profile visualization was performed using R.

### Inorganic compounds quantification

Ammonium and sulfate were quantified with FIA Quikchem 8500 series2 using Quikchem method 10-107-06-2-B Ammonia in surface water, wastewater, and Quikchem method 10-116-10-1-C by FIA (tubidimetric method), respectively. Phosphate was analyzed with FIA Quikchem 8500 series2 with Tecator method ASN 60-01/83 in water by FIA (stannous chloride method). Data acquisition was carried out with Omnion 3.0 from Lachat Instruments. Quantification was performed based on external calibration.

### Data availability

Raw sequencing data are available at Bioproject number PRJNA473500 at http://www.ncbi.nlm.nih.gov/bioproject/.

## Acknowledgements

We are grateful to the Club qualité acéricole Beauce-Appalache for providing the maple samples and to Raymond Nadeau for logistical help and discussions. We are also thankful to the fédération des producteurs acéricoles du Quebec for their financial support.

## Contributions

Study conception and design: MF, CRL, LL, NM

Acquisition of data: NG, MJ, MF, NM

Analysis and interpretation of data: GN, MF

Drafting of manuscript: NG, MF

Critical revision: MF, CRL, NM, MJ, LL, NG

## Footnotes

Supplementary Information accompanies this paper on Journal website

## Supplementary Figure legend

**Supplementary Figure 1.**
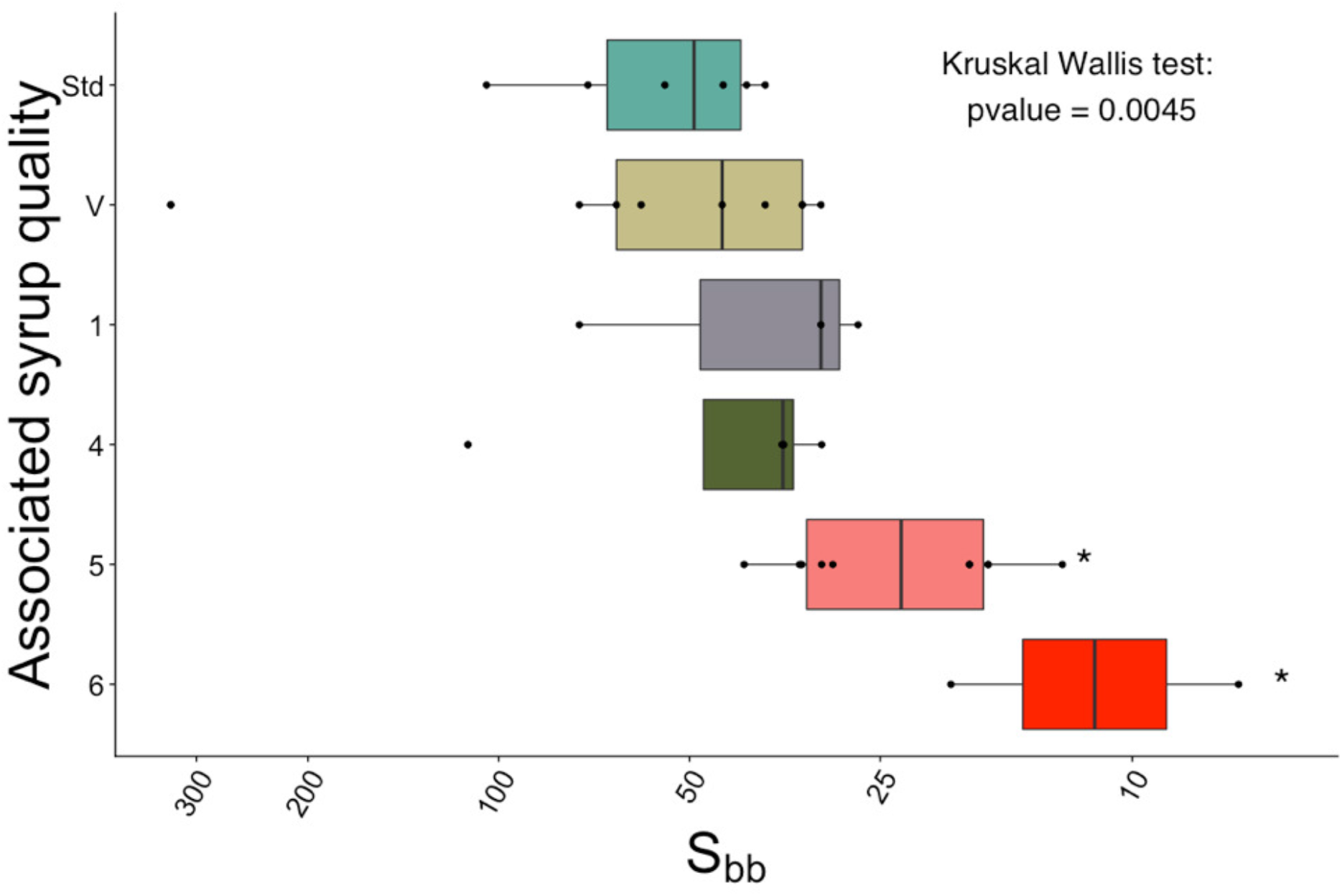
Boxplots representing the sap quality samples from 2016 and 2017 harvesting period displayed by S_bb_. Significant difference is observed between Std - class 5 and Std - class 6 (Dunn’s test P<0.05). Boxplots show the median, 1^st^ and 3^rd^ quantiles.

**Supplementary Figure 2.**
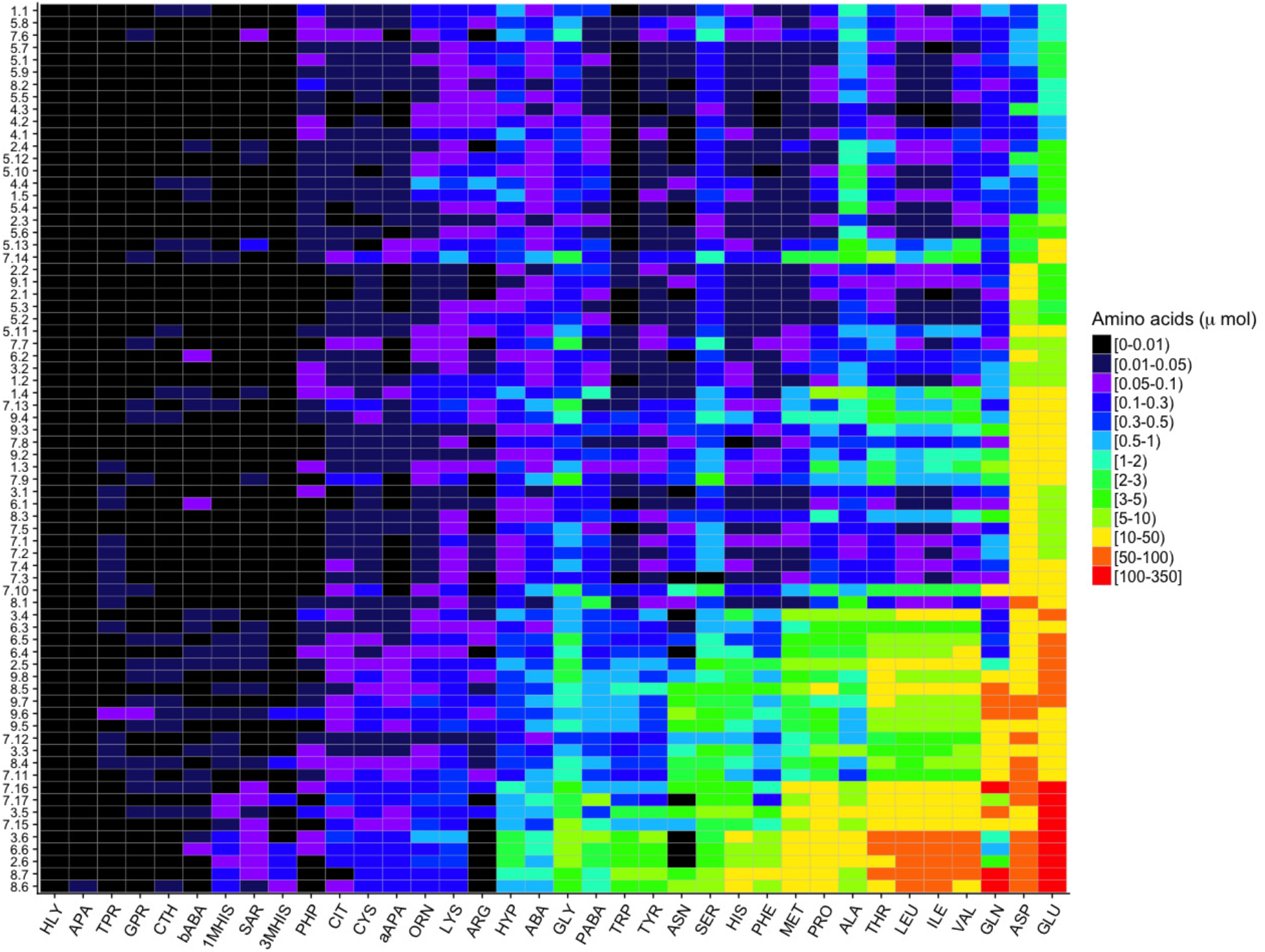
Heatmap of quantified amino acids content in sap samples harvested in 2013. The y-axis displays samples ordered by increasing glutamate concentration. The color scale represents the amino acid concentration in μM.

**Supplementary Figure 3.**
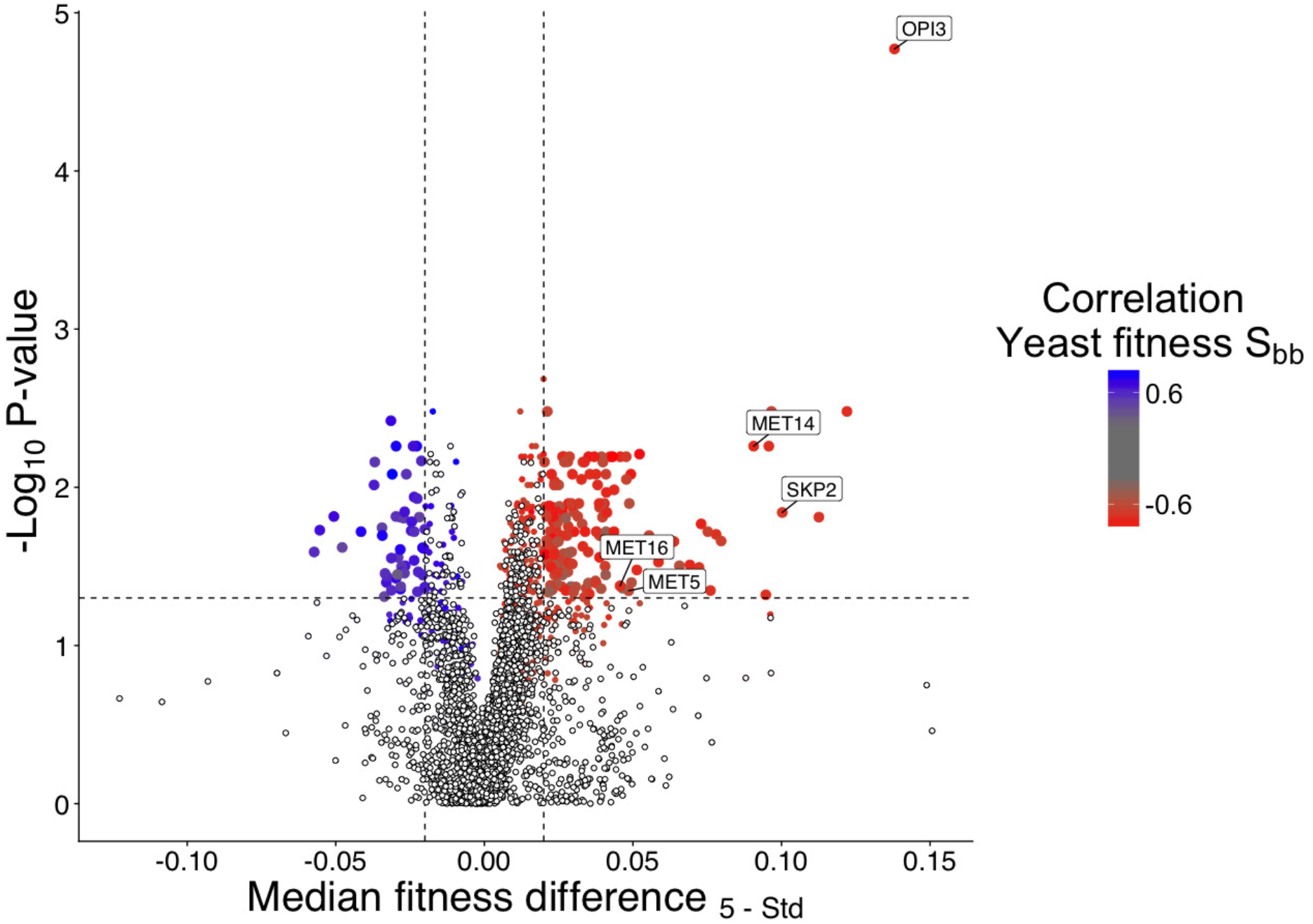
Principal component analysis displaying the sap sample based on the mutant strain’ s fitness from the prototrophic collection. The yeast deletion collection presents a specific fitness profile depending on the sap quality. Std, class 5 and control media are highlighted by dispersal ellipses encompassing 70% of samples.

**Supplementary Figure 4.**
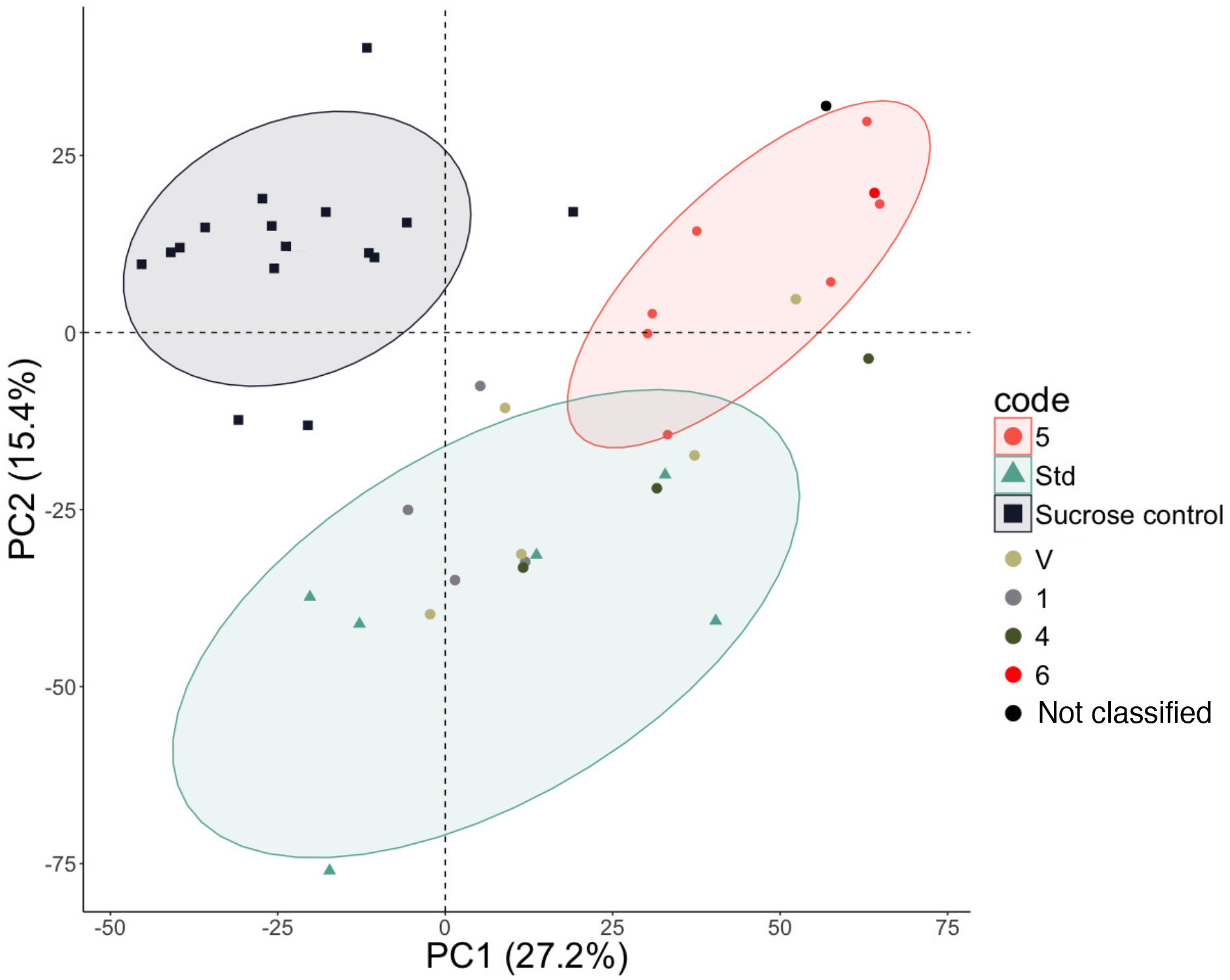
Volcano plot of yeast fitness difference between sap quality (Std vs class 5) with overlaid correlation with S_bb_ as shown by the color scale. For the quality comparison, 218 significant strains were identified by a T-test comparison (adjusted P-value < 0.05 & mean fitness difference +/−0.02). Further, 1035 strains were identified as significatively correlated to the S_bb_ index (Spearman correlation adjusted P-value < 0.01), with 204 strains shared between the two lists. Dots size and color discriminates significant strains for the quality comparison and S_bb_ correlation, respectively. Strains involved in sulfur assimilation are labelled.

**Supplementary Figure 5.**
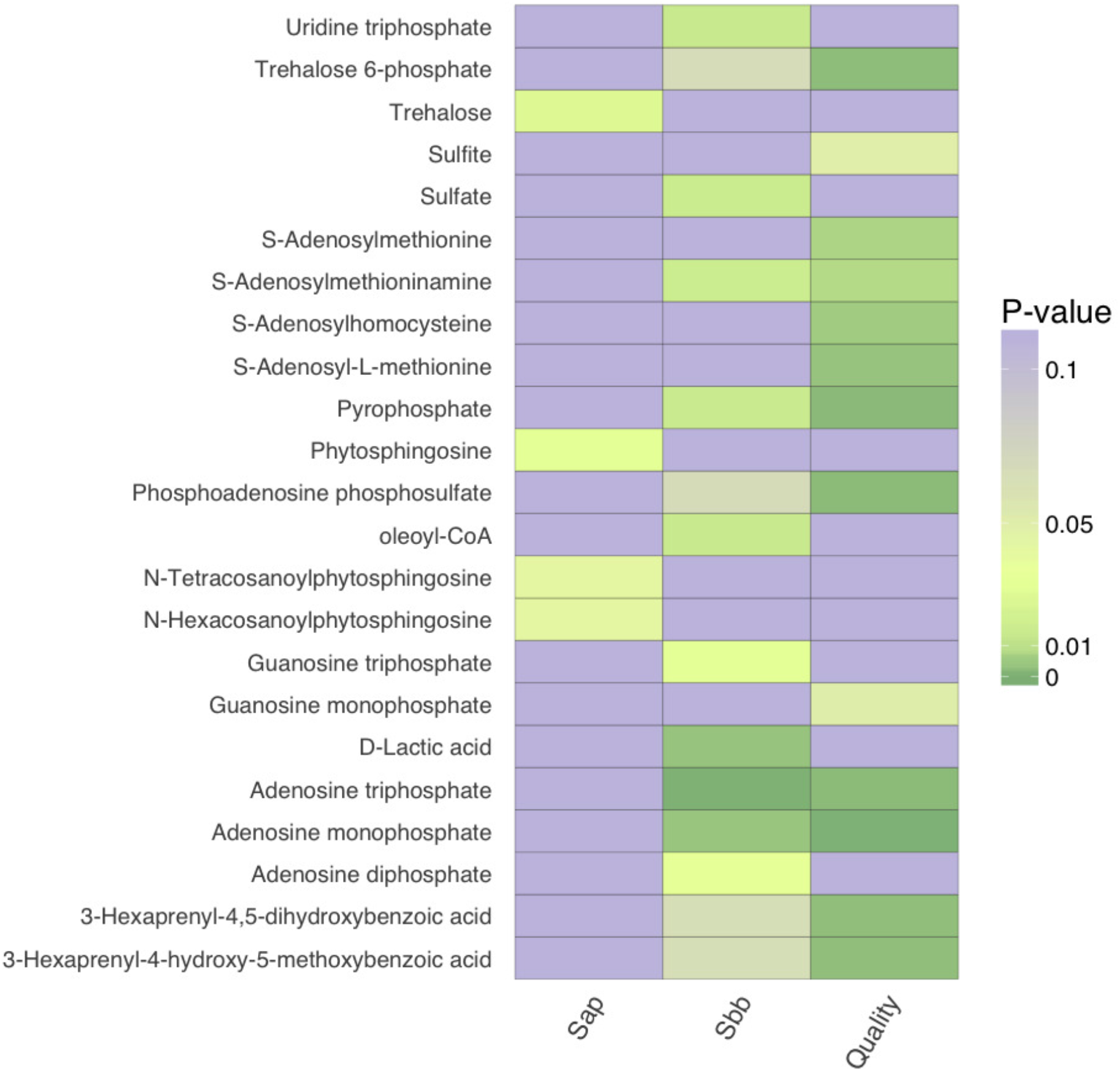
Heatmap of enriched metabolites associated with genes in maple sap, Quality and S_bb_ lists (P-value < 0.01). Gene-metabolites associations were retrieved from YMDB^58^.

